# Differential contributions of low-frequency phase and power in crossmodal temporal prediction: A MEG study

**DOI:** 10.1101/2025.05.26.656128

**Authors:** Rebecca Burke, Jonathan Daume, Till R. Schneider, Andreas K. Engel

## Abstract

**BACKGROUND:** Our representation of time is embedded within multisensory perception, based on sight, sound, or touch. However, despite being a crucial aspect of daily life, the neural dynamics of cross-modal temporal predictions remain elusive. The objective of this study was to investigate neural correlates of tactile-to-visual influences on temporal prediction using magnetencephalography (MEG). We hypothesized to observe increased inter-trial phase consistency (ITPC) in the low-frequency delta range (0.5-4 Hz) due to their involvement in temporal prediction. In addition, stronger ITPC values should correlate with a steeper slope of the psychometric function, indicating phase alignments as a likely cause of more consistent temporal predictions.

**METHODS:** The study was conducted within one MEG session employing a modified version of the time prediction task by Roth et al. [2013] and Daume et al. [2021]. Participants (N=23) observed a visual stimulus moving towards an occluder. Shortly before reaching the occluder, the visual stimulus faded in luminance to make the visual offset less informative. Instead, participants received a brief tactile stimulus to the ipsilateral hand at the time point of disappearance, generating a temporal expectation regarding its reappearance on the opposite side of the occluder. After variable time intervals, a visual stimulus reappeared, and participants had to indicate whether this was *too early* or *too late* compared to the movement before disappearance. A non-predictive control condition involved participants judging the variable luminance of the reappearing visual stimulus compared to its initial luminance at the beginning of the trial. Psychometric curves were fitted to the behavioral data of each participant and condition, and MEG recordings were analyzed using time-frequency representations obtained by wavelet convolution. To compare spectral power and ITPC estimates between conditions within frequency bands showing significant differences to the pre-stimulus baseline, we used cluster-based permutation statistics. Pearson’s correlations were used to examine the relationship between ITPC or power estimates and the steepness of each participant’s psychometric function.

**RESULTS:** ITPC analysis revealed strong increases in the delta range around stimulus disappearance and reappearance. Delta ITPC was significantly stronger during temporal prediction compared to the control condition. Only delta ITPC, but not delta power, correlated with the consistency of temporal prediction. Furthermore, temporal prediction led to increased alpha power in the dorsolateral and medial prefrontal cortex, as well as the right superior temporal sulcus and middle temporal gyrus, whereas beta power did not show differences between conditions.

**CONCLUSION:** Our findings suggest that increased delta ITPC is likely caused by phase resets driven by the temporal prediction process rather than evoked neural activity. Furthermore, our findings indicate that phase alignments occur during crossmodal visuo-tactile-to-visual temporal predictions, even with a combination of non-rhythmic and discrete stimulation. This highlights the broad applicability of phase resets of neural oscillations as a mechanism for predicting timing across various types of stimuli.

## INTRODUCTION

The ability to predict the timing of future events is fundamental to adaptive behavior. The brain continuously generates temporal predictions that allow for the alignment of sensory processing with expected input [Schroeder and Lakatos, 2009, Stefanics et al., 2010, Herbst et al., 2022]. These predictions are thought to be particularly important when sensory input is transiently occluded or ambiguous, enabling the perceptual system to “fill in” missing information and prepare for upcoming stimuli [Summerfield and Egner, 2009]. A fundamental body of research suggests that neuronal oscillations, especially in the delta-(0.5-4 Hz), alpha-(8-12 Hz), and beta- (13-30 Hz) bands, play a central role in temporal prediction [Arnal and Giraud, 2012, Arnal et al., 2015, Calderone et al., 2014, Rohenkohl and Nobre, 2011]. Low-frequency oscillations are ideally suited for encoding temporal regularities due to their alignment with behaviorally relevant timescales [Schroeder and Lakatos, 2009]. In particular, delta-band oscillations have been implicated in phase alignment mechanisms that support sensory anticipation and attentional entrainment [Daume et al., 2021, Burke et al., 2025, Arnal et al., 2015, Cravo et al., 2013]. In the context of rhythmic or temporally structured input, increased inter-trial phase consistency (ITPC) has been interpreted as evidence for phase resetting, whereby the brain aligns ongoing oscillations with anticipated stimulus onsets [Stefanics et al., 2010, Lakatos et al., 2008, Arnal et al., 2015]. While many studies have focused on the auditory modality, there is growing evidence that visual and cross-modal temporal prediction also rely on phase-based neural dynamics [Samaha et al., 2015, van Ede et al., 2011, Daume et al., 2021, Burke et al., 2025]. However, the frequency specificity and topographical distribution of such effects remain debated. Beta-band activity is frequently associated with sensorimotor predictions and has been shown to decrease in response to temporal uncertainty [Daume et al., 2021, Fujioka et al., 2012]. Some studies suggest that visual temporal predictions preferentially modulate alpha-band activity, even when stimuli are presented at delta frequencies. Rohenkohl and Nobre [2011], for example, presented visual stimuli rhythmically at 1.25 and 2.5 Hz, yet only alpha oscillations, rather than delta, showed predictive modulation during stimulus occlusion. This raises the possibility that alpha-band power may play a dominant role in visual prediction, potentially reflecting higher-order cognitive control over sensory anticipation. The neural mechanisms of cross-modal temporal prediction, particularly involving visuo-tactile interactions, remain less well understood. Moreover, few studies have directly compared predictive versus non-predictive tasks using identical sensory stimulation, limiting our understanding of how task-specific cognitive demands shape neural oscillations. Building on the work of Daume et al. [2021], our study introduces a visuo-tactile-to-visual occlusion paradigm that probes temporal prediction across sensory modalities. Further, we employed a control condition, using the exact same sensory stimuli. This design allowed us to isolate the neural effects specific to temporal prediction, independent of sensory input. We focused on two main neural markers: spectral power and ITPC in the delta-, alpha-, and beta-bands, during the critical window of stimulus disappearance. Based on previous work [Daume et al., 2021, Burke et al., 2025], we hypothesized that temporal prediction would be associated with (1) increased delta ITPC reflecting enhanced phase alignment, and (2) reduced beta power indicative of temporal uncertainty. We also explored alpha-band modulations as a potential signature of higher-order cognitive control [Samaha et al., 2015, Rohenkohl and Nobre, 2011]. Furthermore, we tested whether individual differences in ITPC or power correlated with the steepness of the psychometric function, as a measure of temporal prediction precision [Daume et al., 2021]. Our study aims to shed light on the distributed and frequency-specific oscillatory mechanisms supporting temporal prediction in a crossmodal visuo-tactile context.

## MATERIALS AND METHODS

### Participants

Twenty-three healthy adult volunteers participated in the study (9 females; mean age = 26.09 years, SD = 5.06; all right-handed). All participants reported normal or corrected-to-normal vision, no reduced tactile sensitivity, and no history of neurological or psychiatric disorders. Written informed consent was obtained in accordance with the Declaration of Helsinki and the study protocol was approved by the Ethics Committee of the Hamburg Medical Association. Participants received monetary compensation for taking part in the study.

### Experimental procedure

The experimental paradigm comprised two tasks: a temporal prediction (temporal prediction (TP)) task involving visuo-tactile-to-visual temporal prediction, and a luminance matching (luminance matching (LM)) task as a working memory control condition. Both tasks were physically identical in terms of visual and tactile stimulation as well as timing structure but differed in cognitive demands and response criteria. Stimuli were presented on a matte back-projection screen (60 Hz refresh rate; resolution 1920 × 1080 pixels), positioned 65 cm in front of the participant. The main visual stimulus was a small oval (3.5° × 1.0° visual angle), which moved linearly across the screen towards a central white-noise occluder (7.5° × 11.3°), smoothed using a Gaussian filter. The occluder was centered on a gray background (44 cd/m2; RGB = 115), and a red fixation dot was displayed in its center throughout each trial. Participants were instructed to maintain fixation at all times. Each trial began with 1500 ms of fixation. Then, the visual stimulus appeared in the periphery and moved toward the occluder at a speed of 6.9°/s. The onset of stimulus movement was jittered (1000-1500 ms before disappearance) to prevent temporal predictability and the initial side of stimulus onset (left or right) was counterbalanced across participants but remained constant within individuals. Importantly, the visual stimulus decreased in luminance, thereby making the visual offset less informative. Instead, participants received a brief tactile cue (70 ms) at the time point of full disappearance to the index finger of the side where the stimulus was moving toward the occluder. The occluder size and stimulus speed were calibrated such that a constant time of 1500 ms was required to move behind the occluder. After disappearance, the stimulus reappeared at variable time points (jittered from *±*17 ms to *±*467 ms around the expected reappearance at 1500 ms), resulting in 20 different reappearance latencies. This temporal manipulation allowed assessment of subjective timing perception. The initial luminance of the moving stimulus changed on a trial-by-trial level. Similarly, the luminance changed after reappearance relative to the initial luminance (range: *±*1 to *±*40 cd/m2; in 20 steps). These values were matched across conditions to ensure the same physical stimulus structure.

In the TP task, participants had to judge whether the reappearing stimulus appeared either *too early* or *too late*, based on the previously perceived visual motion before occlusion. The tactile stimulus was administered using a Braille piezostimulator (QuaeroSys, Stuttgart, Germany; 24 pins, 1 mm diameter, 2.5 mm spacing), activating all pins simultaneously. At the time point of tactile stimulation, no concurrent visual event occurred on the screen. Responses were given via button press using the hand contralateral to the stimulus reappearance side. Response mapping (*too early* = left key / *too late* = right key, or vice versa) was counterbalanced across participants. In the LM control task, participants were instructed to judge whether the luminance of the reappearing stimulus was *brighter* or *darker* than the pre-disappearance luminance. All other aspects of the physical stimulation and timing were identical to the TP task. This task served as a cognitive control to account for non-temporal processing demands and ensured identical visual stimulation between tasks. The experiment was conducted in a single recording session, including 12 blocks (6 per condition), resulting in 60 trials per block and a total of 720 trials per participant (360 per condition). Each participant completed a brief initial practice of the task, including feedback after each trial. For the main experiment, participants received feedback at the end of each block in which the overall accuracy was displayed. Participants could rest as needed after each block. To mask the sound of the Braille stimulator, participants wore in-ear MEG-compatible headphones delivering continuous pink noise (85 dB, 48 kHz sampling rate) during all experimental blocks.

### Data acquisition and preprocessing

The electrophysiological data was recorded with a 275-channel whole-head MEG system (CTF MEG International Services LP, Coquitlam, Canada) at 1200 Hz in a dim, magnetically shielded room. Electrooculogram (EOG), electromyogram (EMG), and electrocardiogram (ECG) were recorded using Ag/AgCl electrodes for artifact rejection. Head position was monitored online and realigned before each block if deviations exceeded 5 mm from the original position [Stolk et al., 2013]. We used MATLAB (R2016b) (The MathWorks Inc., 2019), FieldTrip [Oostenveld et al., 2011], and custom scripts to analyze the data. Preprocessing included bandpass filtering (0.5-170 Hz), notch filters at 50, 100, and 150 Hz and downsampling the data to 400 Hz. Trials with artifacts (e.g., muscle, jumps) were excluded via a semi-automatic rejection procedure. On average, 666.6 *±*37.4 (92.58% *±*5.14%) trials remained after preprocessing. Finally, independent component analysis (ICA) was used to remove ocular, cardiac, and muscle artifacts. Approximately 25 *±*6.9 out of 275 components were removed per participant. Furthermore, trials deviating by ≥1 frame (17 ms) from the intended stimulus timing were excluded. Sensors were flipped to ensure comparability across participants who viewed stimuli from different directions (see Time-frequency analysis).

### Behavioral analysis

Reaction times (RT) and accuracy were analyzed using R (R Core Team, 2024) and RStudio (RStudio Team 2020). Psychometric curves were calculated in MATLAB (The MathWorks Inc., 2019). Trials with reaction times >3 SDs from the mean were excluded. RTs were logarithmically transformed and standardized prior to statistical modeling. Psychometric curves were fitted using binomial logistic regression (MATLAB’s *glmfit*.*m* and *glmval*.*m*) to determine each participant’s subjective point of right on time (ROT) in the prediction task and point of subjective equality (PSE) in the working memory control task. The steepness of the psychometric function was quantified using the inverse difference between 25% and 75% threshold points. One-sample *t*-tests (*α* = 0.025, Bonferroni-corrected) were used for comparisons of the two tasks.

### Time-frequency analysis

Time-frequency decomposition was performed via convolution with 40 complex Morlet wavelets (0.5-100 Hz). Spectral power was computed for each trial and normalized to a pre-stimulus baseline (−500 to -200 ms). Power estimates were binned into 100 ms steps and averaged per condition. Trials were segmented into four overlapping time windows: Baseline: -550 to -50 ms before stimulus motion onset; Movement: -50 to 950 ms from motion onset; Disappearance: -350 to 950 ms from full occlusion; Reappearance: -350 to 450 ms from stimulus reappearance; For statistical comparisons between tasks, we used cluster-based permutation tests [Maris and Oostenveld, 2007], correcting for multiple comparisons across time, frequency, and space. Sensor data were flipped along the sagittal axis for participants who viewed leftward motion.

Our analyses focused primarily on the Disappearance window, since this is thought to be the critical time window for the temporal prediction process. Furthermore, for source localization we used Dynamic Imaging of Coherent Sources (DICS) beamforming [Gross et al., 2001] based on individual magnetic resonance images and a single-shell volume conductor model [Nolte, 2003] with a 5003 voxel grid aligned to the MNI152 template brain. Again, as for sensor-level analysis, all voxels were flipped along the sagittal axis for participants who were presented with the white ellipse moving from right to left prior to statistical calculations.

### Inter-trial phase consistency (ITPC)

ITPC was computed from complex time-frequency representations obtained via wavelet convolution (see above). Phase angles were extracted at each time point and trial using MATLAB’s *angle*.*m* function, and ITPC was calculated as:

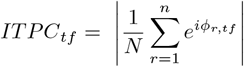

where *n* is the number of trials and *ϕ*_*r,tf*_ the phase angle in trial *r* at time frequency point *tf*. ITPC values were averaged in 100 ms bins and pooled across channels and conditions to provide an overview of phase alignment throughout the trials. Furthermore, to focus on dynamics surrounding stimulus disappearance, we additionally computed ITPC in a -1,900 to 1,900 ms window centered on full occlusion of the white ellipse. Values were averaged within the 0.5-4 Hz delta-band and compared between conditions using cluster-based permutation statistics over time bins and sensors. At the source level, ITPC was estimated using the same beamformer filters as in the spectral power analysis. Finally, to assess neural-behavioral coupling, voxel-wise Pearson correlations were performed between ITPC (averaged 0 to 800 ms post-disappearance) and the steepness of each participant’s psychometric function. Multiple comparisons were corrected via cluster-based permutation testing.

## RESULTS

### Comparable behavioral performance across conditions

To assess behavioral differences between the TP and LM conditions, we compared behavioral accuracy, reaction times, as well as the slope and bias of the psychometric curve. Accuracy was significantly lower in the TP condition (mean = 0.719, SEM = 0.060) compared to LM (mean = 0.764, SEM = 0.034) (*t*(22) = -4.21, *p* < .001, Cohen’s *d* = -0.88). Therefore, participants performed more accurately when judging luminance than when making temporal predictions. However, a more sensitive measure of performance, the psychometric curve, showed no statistically significant difference in the slope of responses between the TP and LM conditions (*t*(22) = -1.89, *p* = .072, Cohen’s *d* = -0.39). Similarly, the bias did not differ significantly between conditions (*t*(22) = -0.30, *p* = .764, Cohen’s *d* = -0.06). These results show comparable response trends and discrimination sensitivity between conditions. Analysis of log-transformed and standardized reaction times showed no significant overall difference between conditions (*t*(22) = -0.59, *p* = .559, Cohen’s *d* = -0.12).

### Alpha power significantly increased during temporal prediction

We first examined time-resolved total power across frequencies, averaged over all conditions, sensors, and participants. To characterize general oscillatory dynamics during the trial, we applied cluster-based permutation statistics to compare the TP and LM task against baseline (Figure 2). Relative to the pre-stimulus baseline, a significant increase in delta-band power was observed during the disappearance window, time-locked to the stimulus occlusion (all cluster-*p*<.001). In contrast, beta-band power showed a significant decrease following stimulus disappearance, consistent with previous reports of beta suppression during prediction processes. Similarly, we found significant alpha-band modulations during the occlusion window. To isolate condition-specific effects, cluster-based permutation statistics were applied to compare the TP and LM tasks against each other within the delta-, alpha-, and beta-band on sensor and source level (Figure 2B&C). While no significant clusters were found for delta or beta power, we observed a significant increase in alpha-band power during the TP task compared to the LM task in the dorsolateral and medial prefrontal cortex. A second cluster was found in the area of middle temporal gyrus and superior temporal sulcus (both cluster-*p*=.02).

**Figure 1:**
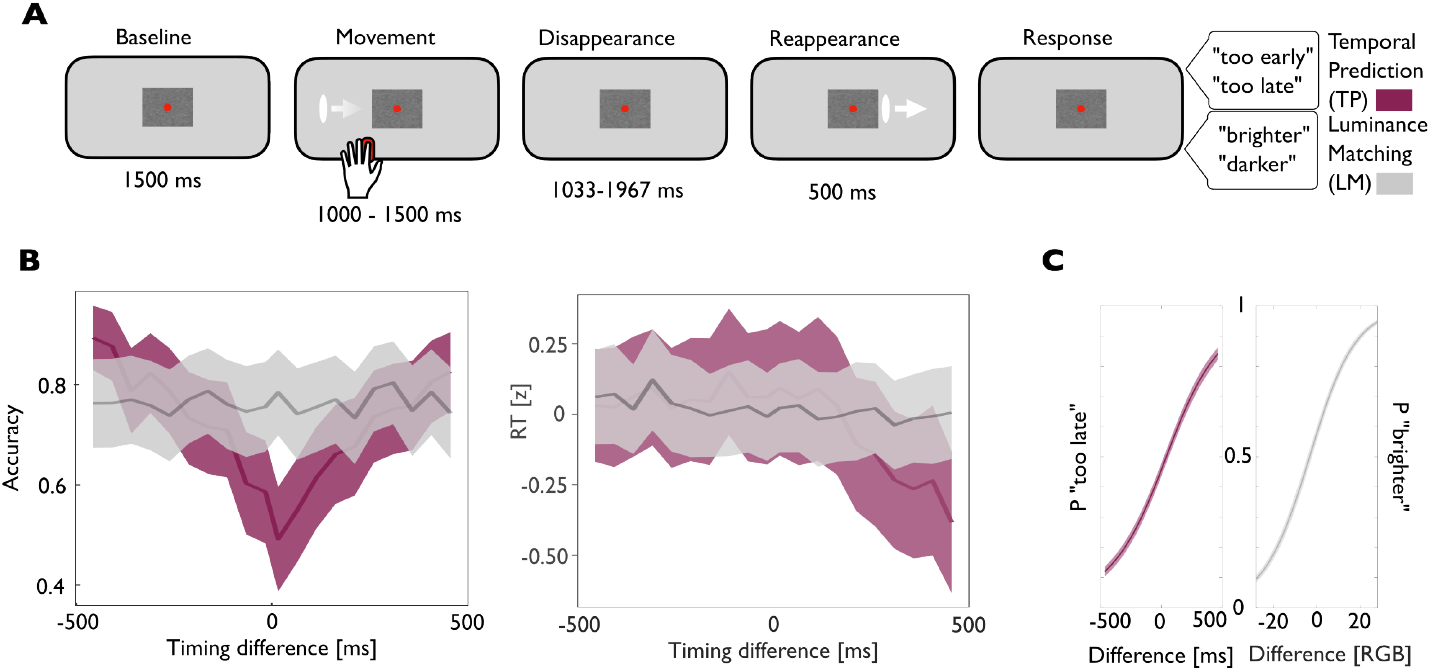
Experimental design and behavioral results. (A) Experimental paradigm. Participants viewed a white ellipse moving toward and disappearing behind an occluder. Just before disappearance, the ellipse faded in contrast to make the visual offset less informative. Instead, participants received a brief tactile cue to the ipsilateral hand at the time of disappearance. Therefore, the timing of the tactile stimulus provided an important cue needed to judge the appropriateness of the reappearance of the visual stimulus. After a variable delay, the ellipse reappeared with a slightly different luminance. In the temporal prediction (TP) task, participants judged whether the reappearance was *too early* or *too late*, based on the velocity of the stimulus but ignoring the luminance change. In the luminance matching (LM) control task, visual and tactile stimulation identical to the TP task was used, but participants judged the brightness of the reappearing stimulus, ignoring its timing. (B) Accuracy and reaction times along timing differences. The left panel shows participant accuracy across various timing differences, with time 0 ms indicating the objectively correct reappearance after 1500 ms for the TP (red) and LM (grey) condition. The right panel displays log-transformed and standardized reaction times (RT) across timing differences. (C) Psychometric curves illustrate response sensitivity and bias in the TP (red) and LM (grey) condition. Time 0 ms in the TP condition (left) refers to the objectively correct reappearance, while a luminance difference of 0 RGB values in the LM condition (right) indicates objective luminance equality upon reappearance. *P*=proportion, RGB=red-green-blue.

**Figure 2:**
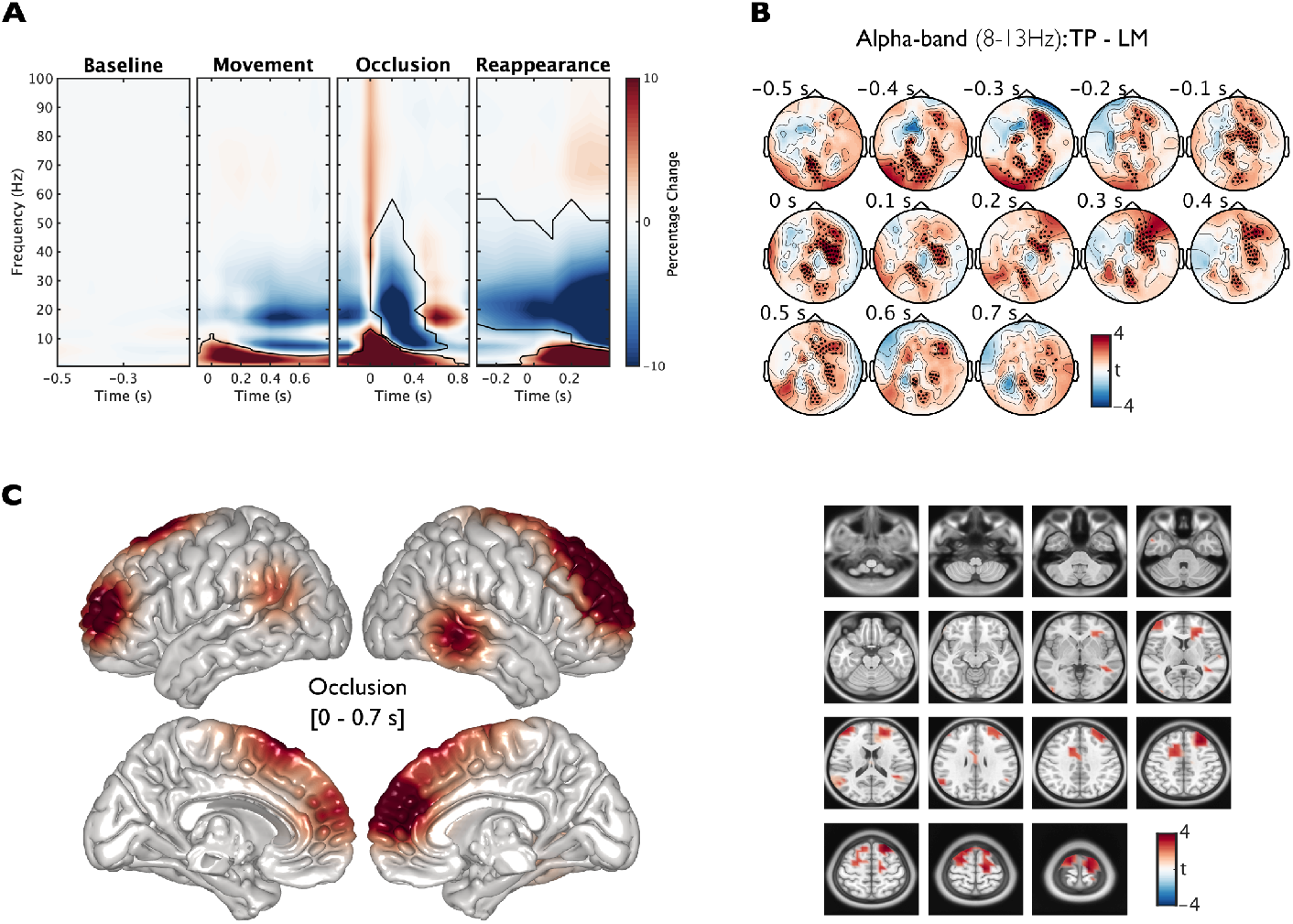
Spectral power modulations. (A) Spectral power was averaged across all sensors, conditions, and participants, with each time window aligned to key events in the taks and normalized using a pre-stimulus baseline. Time 0 s marks the onset of each event. Significant power changes relative to baseline are indicated by areas enclosed in solid lines, as determined by cluster-based permutation testing. (B, C) Comparison of alpha-band activity (8–13 Hz) between the temporal prediction and luminance matching task during time intervals surrounding stimulus occlusion. (B) On sensor level, black dots mark sensors within clusters that showed statistically significant differences between conditions. (C) At the source level, cluster-based permutation tests identified voxel clusters with significant task-related differences, visualized as colored regions in the surface and slice images averaged across the disappearance time window (0-0.7s).

### Delta inter-trial phase consistency was increased during and correlated with temporal prediction

Similarly to our spectral power analysis, we examined time-resolved ITPC across frequencies, averaged across all conditions, sensors, and participants (Figure 3A). Cluster-based permutation statistics revealed a significant ITPC increase in all frequency bands at the time point of disappearance compared to pre-stimulus baseline (all cluster-*p*=<.001). To assess task-specific differences, we compared the TP and LM tasks using cluster-based permutation statistics. This revealed significantly stronger ITPC in the delta-band during the disappearance window for the TP task compared to the LM task (cluster-*p*=.02). This effect was most pronounced within the time window from 0 to 800 ms following stimulus disappearance, indicating enhanced phase alignment across trials when participants were engaged in predicting the reappearance of the visual stimulus (Figure 3B). Source-level analysis of this contrast revealed increased delta-band ITPC in several visual and visual-association regions, including the left and right primary and secondary visual cortices as well as the right inferotemporal cortex (cluster-*p*=.01). The pattern of increased phase consistency in early visual areas, particularly contralateral to the anticipated stimulus, supports the interpretation of a phase reset mechanism in response to internally generated temporal predictions. Finally, correlational analyses revealed a significant correlation between delta ITPC and the steepness of the individual psychometric function, with clusters primarily located in the bilateral primary somatosensory cortex and sensory association areas (Figure 4) (cluster-*p*=.03). No such correlation was observed for delta power, nor did alpha power correlate significantly with psychometric slope.

**Figure 3:**
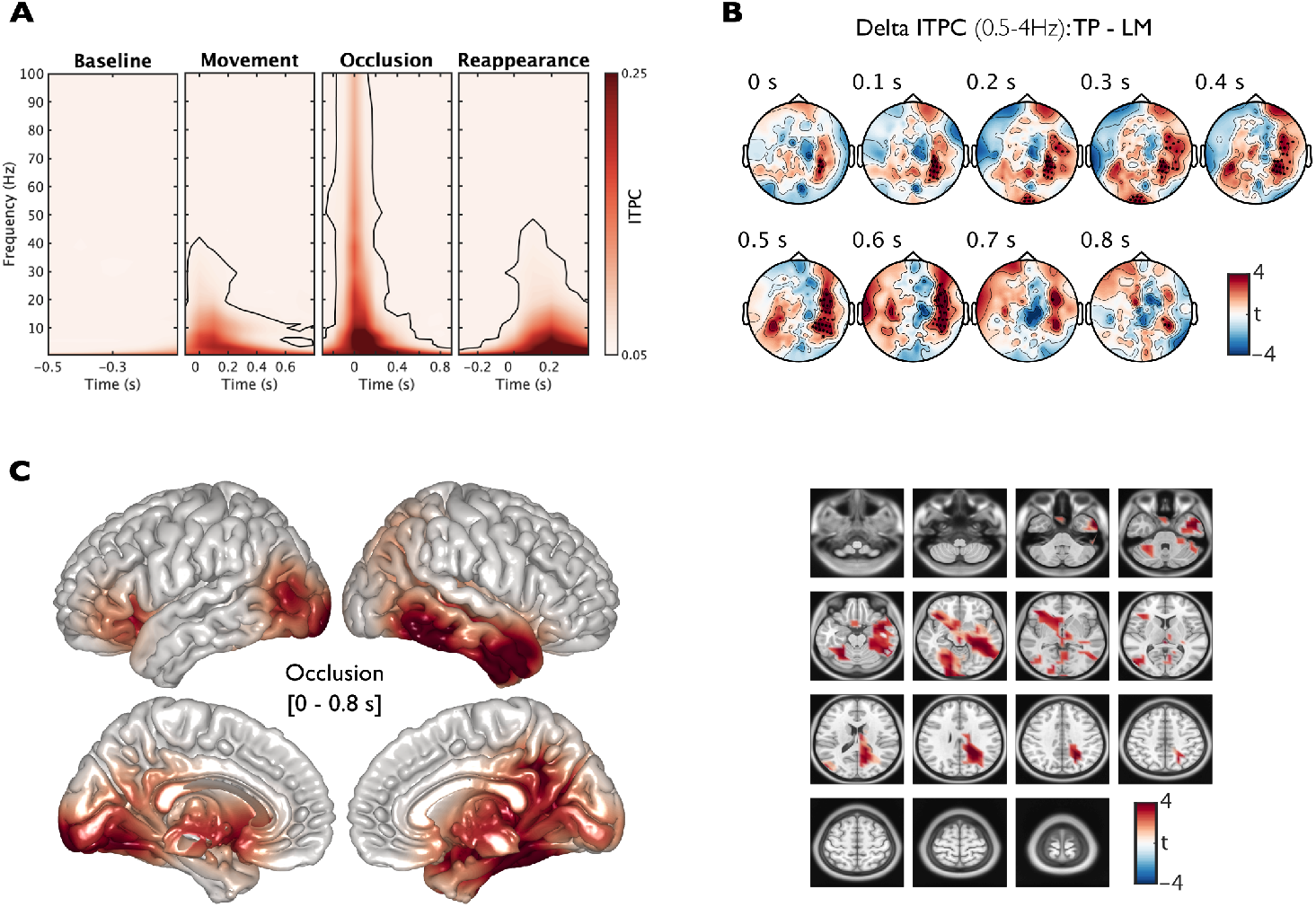
ITPC. (A) ITPC values were averaged across all sensors, conditions, and participants, with significant deviations from baseline highlighted by outlines derived from cluster-based permutation testing. (B, C) Differences in delta-band ITPC (0.5–3 Hz) between the temporal prediction task and the luminance matching control. (B) Black dots mark sensors that formed significant clusters in the group comparison on sensor level. (C) On source level, colored regions indicate voxel clusters with statistically significant differences between conditions.

**Figure 4:**
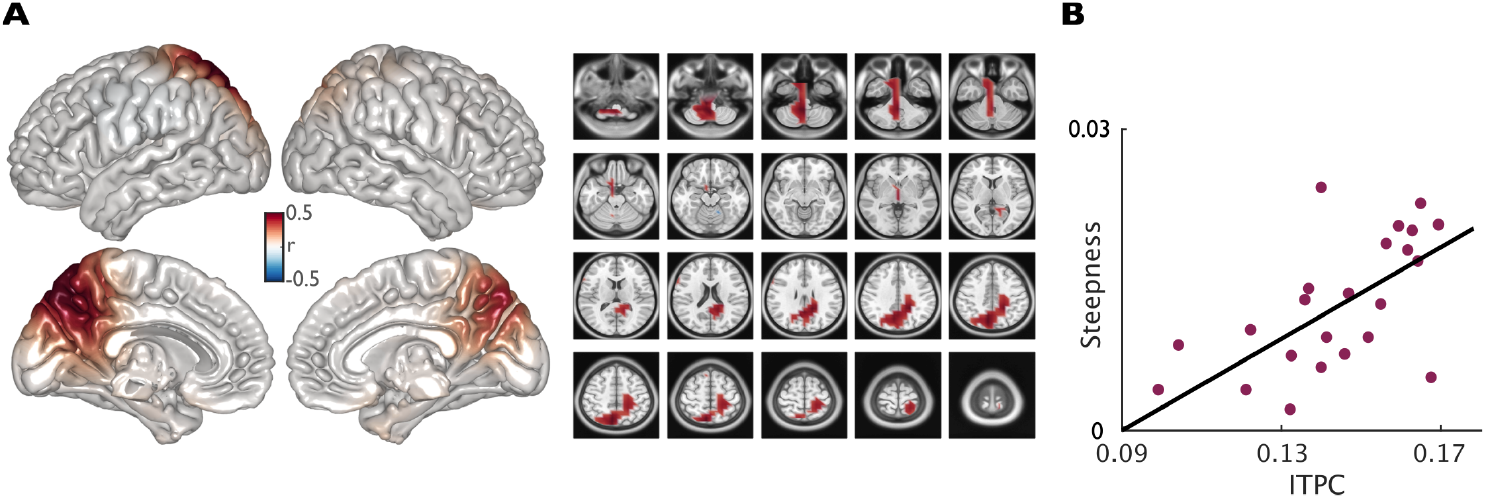
Correlation of delta ITPC and behavior. (A) Voxel-wise correlations between individual ITPC values and the steepness of participants’ psychometric functions are shown for the temporal prediction task. ITPC was computed in the delta-band (0.5–4 Hz) over a time window from 0 to 800 ms following stimulus disappearance (see Figure 3). Colored regions highlight voxel clusters where correlations reached statistical significance using cluster-based permutation statistics. (B) The scatter plot shows this correlation, with each dot representing one participant; ITPC values were averaged across voxels within the significant clusters. No significant correlations were found for the luminance matching condition or between spectral power in the delta-, alpha-, and beta-band and behavioral measures.

## DISCUSSION

The current study investigated neural oscillatory dynamics during a visuo-tactile-to-visual temporal prediction task, with a luminance matching task as a cognitive control.

### Beta desynchronization during visual occlusion occurs across predictive and non-predictive tasks

In line with previous studies, we observed a significant reduction in beta-band power during the disappearance window, relative to pre-stimulus baseline. This suppression occurred in both the TP and LM conditions, and while the effect was robust, no significant task difference emerged. Beta power suppression could reflect processes engaged during both predictive and non-predictive evaluation of dynamic sensory events. Beta-band oscillations, particularly over sensorimotor and associative cortices, have been widely implicated in top-down predictive control, timing, and maintenance of the current sensorimotor set [Engel and Fries, 2010, Arnal and Giraud, 2012, Fujioka et al., 2012, Kilavik et al., 2013]. The transient suppression of beta power in our task is consistent with the notion that beta desynchronization facilitates adaptive updating of internal models in response to sensory transitions. In our study, this would occur due to the sudden occlusion of the visual stimulus. Importantly, beta suppression has also been linked to increased temporal uncertainty, with reduced beta power observed when stimulus timing becomes unpredictable or requires estimation [Arnal et al., 2015, Palmer et al., 2019, Todorovic et al., 2015]. However, the absence of a task-specific difference in beta power between TP and LM conditions is notable. Given that both tasks involved identical sensory transitions and attentional demands, but only TP required active temporal estimation, it is possible that beta-band dynamics encode uncertainty in a modality- or context-unspecific manner. For example, beta suppression may signal increased reliance on internal models during periods of occlusion, regardless of whether a specific temporal prediction is required, thus serving as a non-specific readiness or preparatory mechanism across tasks [Palmer et al., 2019].

### Increased alpha power during temporal prediction suggests top-down engagement

Alpha oscillations constitute a central component of the neural architecture underpinning temporal prediction in multisensory contexts [Keil and Senkowski, 2018]. Our results are in line with this by showing that alpha-band power increased significantly during the TP compared to LM condition in a cluster of voxels in the dosolateral prefrontal cortex (DLPFC) and medial prefrontal cortex (mPFC) (BA 9), despite both tasks involving identical sensory input. A second alpha cluster emerged in middle temporal gyrus (MTG)/superior temporal sulcus (STS), regions typically implicated in timing and multisensory integration [Beauchamp et al., 2008, Noesselt et al., 2012]. These regions appear to leverage dynamic modulations in alpha-band activity to implement top-down expectations. Specifically, event-related alpha power increases, are believed to transiently inhibit task-irrelevant sensory processing, in line with the gating-by-inhibition framework [Jensen and Mazaheri, 2010, Klimesch et al., 2007]. In this framework, enhanced alpha power does not reflect a passive state, but an active process of internal attentional control, aimed at suppressing potentially interfering sensory input during task phases that rely on internal computation rather than external stimulation. The prefrontal cortex (PFC), specifically the DLPFC and mPFC, are repeatedly implicated in orchestrating these forms of internal control. During the occlusion phase of our task, when no sensory input is present and participants must internally maintain a dynamic temporal model to judge the timing of the stimulus reappearance, increased alpha power in frontal regions likely reflect inhibition of premature or irrelevant processing (e.g., of sensory noise or competing expectations) [Klimesch et al., 2007, Sauseng et al., 2005]. Meanwhile, temporal regions including the STS and MTG have been implicated in adjudicating the temporal alignment of incoming multisensory stimuli, modulating alpha power in accordance with the temporal congruency or asynchrony of visuo-tactile inputs [Noesselt et al., 2012]. In an fMRI study, Noesselt et al. [2012] examined how the brain encodes synchronous vs. asynchronous audiovisual stimuli. They identified subregions along the STS that preferentially responded when stimuli were out of sync versus in sync. Notably, when participants perceived asynchrony, there was stronger functional coupling between the STS complex and prefrontal areas. Therefore, these regions may be inherently connected during temporal processing tasks. Although our temporal prediction task is contrasted against a working memory control condition, it is important to recognize that temporal prediction itself recruits executive processes analogous to those used in working memory, including the maintenance and updating of internal representations over time. Specifically, during the occlusion period, participants must hold in mind the velocity and trajectory of a moving stimulus and project it forward to estimate the expected time of reappearance, a process that closely parallels processes in visuospatial working memory tasks [Sauseng et al., 2005]. While both tasks rely on internal processing and top-down control, the working memory task involves the maintenance of static visuospatial information, with comparatively less emphasis on internal modeling over time, which may enhance alpha synchronization due to increased task demands [Klatt et al., 2022]. In summary, the observed increase in alpha power in the TP condition, relative to the LM condition, supports the view that alpha-band oscillations play a central role in implementing top-down control during internally guided temporal prediction processes. The engagement of both prefrontal and temporal regions suggests a coordinated network in which increased alpha synchronization supports the maintenance of temporal expectations by actively suppressing premature or distracting input. These findings may extend existing models of alpha-based inhibition by demonstrating that such mechanisms are not only recruited during classical working memory or attention tasks but also during anticipatory processes. Thus, alpha oscillations could reflect a dynamic control process that is flexibly deployed depending on the temporal structure and cognitive demands of a task, reinforcing their role as a fundamental neural mechanism for predictive processing in multisensory contexts.

### Delta phase alignment may support crossmodal predictive timing

A central finding of our study is the enhanced delta-band ITPC during stimulus disappearance in the TP condition compared to the LM control condition. This supports the idea that delta oscillations subserve temporal prediction by providing a phase-based reference frame for aligning internal neural states with anticipated external events [Arnal and Giraud, 2012, Schroeder and Lakatos, 2009, Cravo et al., 2013]. The increase in ITPC was most pronounced between 0 and 800 ms after disappearance of the stimulus, corresponding to the time window in which participants internally simulated the continuation of the visual trajectory. Our results directly replicate the findings of Daume et al. [2021], who reported increased delta-band ITPC during a unimodal visual and crossmodal visuo-tactile temporal prediction task. While our findings conceptually align with the work by Daume et al. [2021], the visuo-tactile-to-visual paradigm introduces a novel and more demanding form of crossmodal temporal prediction. The previous design assessed whether visual input could prime the somatosensory system to anticipate tactile onset. In contrast, the novel task reverses this logic: we presented a moving visual stimulus that gradually decreased in luminance prior to disappearing, thereby rendering the visual offset temporally ambiguous. Crucially, just before disappearance, a brief, tactile stimulus was delivered to the index finger. This tactile cue served as a precise temporal anchor for participants to initiate an internal prediction about the timing of a subsequent visual reappearance from behind the occluder. This subtle yet important change in task structure entails a crossmodal shift in predictive responsibility: the tactile system now supplies the timing cue, while the visual system becomes the target of temporal anticipation, necessitating participants to integrate temporally informative, non-redundant cues across modalities. Our results thus extend the phase reset framework by demonstrating that delta-band ITPC tracks crossmodal predictions in contexts where temporal structure is not visually salient, but is conveyed through a secondary modality. This supports the interpretation that delta oscillations facilitate the binding of temporally disjoint, crossmodal cues into coherent predictive representations, even when sensory input is ambiguous or degraded. Importantly, delta ITPC was positively correlated with individual differences in temporal prediction performance, providing further evidence that phase alignment is not only a marker of neural synchronization but also reflects behaviorally meaningful anticipatory processes. Source localization revealed increased delta ITPC in bilateral early visual cortices (V1/V2), consistent with research showing that primary and secondary occipital areas exhibit phase alignment during visual anticipation [Cravo et al., 2013]. Additionally, robust delta-phase synchronization was observed in right-lateralized inferotemporal cortex, a region implicated in high-level visual processing and predictive coding of object identity and motion trajectories [Summerfield and Egner, 2009]. Importantly, our findings also implicate regions beyond the sensory cortices. Enhanced delta ITPC in the cerebellum aligns with extensive literature attributing to the cerebellum a key role in subsecond timing and sensorimotor prediction [Ivry and Keele, 1989, Merchant et al., 2013]. This supports the view that cortico-cerebellar loops contribute to temporal anticipation by modulating the timing of cortical excitability through slow oscillatory dynamics. Notably, we also found that delta ITPC in bilateral somatosensory and sensory association cortices significantly correlated with the steepness of individual psychometric functions, indicating a direct relationship between neural phase alignment and the precision of temporal predictions. These results support earlier findings showing delta-phase consistency contributes to timing accuracy and crossmodal temporal integration [Breska and Ivry, 2017, Cravo et al., 2013]. The absence of corresponding correlations with delta power underscores the specific functional relevance of phase-based measures in capturing behaviorally relevant neural dynamics [Haegens and Golumbic, 2018]. Together, these results highlight delta ITPC as a robust index of temporal prediction across a distributed network encompassing sensory, associative, and subcortical structures, with functionally specific contributions to crossmodal timing precision and anticipatory behavior.

## CONCLUSION

These results contribute to a growing literature, suggesting delta oscillations as a scaffold for predictive timing, especially under uncertainty. The lack of task-specific effects in beta power, and the task-dependent modulation of alpha activity in frontal and temporal regions further suggest that temporal prediction recruits a flexible hierarchy of oscillatory mechanisms at multiple timescales [Senkowski and Engel, 2024]. Importantly, this data underscores the need for studies using causal manipulations, such as phase-specific transcranial alternating current stimulation (tACS), to determine whether phase alignment is not only correlated with, but causally drives, enhanced temporal prediction.

## Data availability statement

Data to evaluate the results will be made available upon reasonable request to the authors.

## CRediT authorship contribution statement

**Rebecca Burke:** Formal analysis, Validation, Visualization, Data curation, Writing - original draft, Writing - review & editing. **Jonathan Daume**: Conceptualization, Methodology, Investigation, Writing - review & editing. **Till R. Schneider**: Conceptualization, Methodology, Writing - review & editing, Supervision. **Andreas K. Engel**: Conceptualization, Methodology, Writing - review & editing, Funding acquisition, Project administration, Supervision.

## Acknowledgements

This work was supported by grants from the Deutsche Forschungsgemeinschaft (SFB TRR 169/B1/B4 awarded to A.K.E.) and from the European Research Council (project cICMs, ERC-2022-AdG-101097402 awarded to A.K.E.). We thank Christiane Reißmann for assistance in data recording, and Alexander Maÿe, Marleen J. Schoenfeld, and Peng Wang for helpful discussions on the data. Views and opinions expressed in this paper are those of the authors only and do not necessarily reflect those of the European Union or the European Research Council. Neither the European Union nor the granting authority can be held responsible for them.

